# Pollinator size and its consequences: Predictive allometry for pollinating insects

**DOI:** 10.1101/397604

**Authors:** Liam K. Kendall, Romina Rader, Vesna Gagic, Daniel P. Cariveau, Matthias Albrecht, Katherine C. R. Baldock, Breno M. Freitas, Mark Hall, Andrea Holzschuh, Francisco P. Molina, Joanne M. Morten, Janaely S. Pereira, Zachary M. Portman, Stuart P. M. Roberts, Juanita Rodriguez, Laura Russo, Louis Sutter, Nicolas J. Vereecken, Ignasi Bartomeus

## Abstract

1. Body size is an integral functional trait that underlies pollination-related ecological processes, yet it is often impractical to measure directly. Allometric scaling laws have been used to overcome this problem. However, most existing models rely upon small sample sizes, geographically restricted sampling and have limited applicability for non-bee taxa. Predictive allometric models that consider biogeography, phylogenetic relatedness and intraspecific variation are urgently required to ensure greater accuracy.
2. Here, we measured body size, as dry weight, and intertegular distance (ITD) of 391 bee species (4035 specimens) and 103 hoverfly species (399 specimens) across four biogeographic regions: Australia, Europe, North America and South America. We updated existing models within a Bayesian mixed-model framework to test the power of ITD to predict interspecific variation in pollinator dry weight in interaction with different co-variates: phylogeny or taxonomy, sexual dimorphism and biogeographic region. In addition, we used ordinary least squares (OLS) regression to assess intraspecific dry weight – ITD relationships for 10 bee and five hoverfly species.
3. Including co-variates led to more robust interspecific body size predictions for both bees (Bayesian *R*^2^: 0.946; Δ*R*^2^ 0.047) and hoverflies (Bayesian *R*^2^: 0.821; Δ*R*^2^ 0.058) relative to models with ITD alone. In contrast, at the intraspecific level, our results demonstrate that ITD is an inconsistent predictor of body size for bees (*R*^2^: 0.02 – 0.66) and hoverflies (*R*^2^: −0.11 – 0.44).
4. Therefore, predictive allometry is more suitable for interspecific comparative analyses than assessing intraspecific variation. Collectively, these models form the basis of the dynamic *R* package, ‘*pollimetry*’, which provides a comprehensive resource for allometric research concerning insect pollinators worldwide.

## Introduction

Body size is an important functional trait that influences ecological patterns across all levels of biological organisation. In insects, adult body size variation is the outcome of natural selection affecting physiological and biochemical processes during ontogeny (Chown & Gaston 2010). For example, body size impacts metabolic and growth rates (Angilletta et al. 2004; Ehnes et al. 2011), life history (e.g. life span and reproductive rate; Speakman 2005; Teder et al. 2008) and ecological attributes, such as species abundance, trophic interactions, geographic range size and dispersal ability (Brown et al. 2004; White et al. 2007; Stevens et al. 2012; Velghe & Gregory-Eaves 2013; DeLong et al. 2015). In addition, body size can drive key ecosystem functions and services such as decomposition, carbon cycling, predation, primary productivity and pollination (Woodward & Hildrew 2002; Greenleaf et al. 2007; Rudolf & Rasmussen 2013; Garibaldi et al. 2015; Schramski et al. 2015).

Body size is most commonly measured as specimen dry weight. As such, obtaining direct measurements can be impractical and time consuming. Direct measurements often require destructive methods, which is unfavourable for museum specimens and threatened species (Rogers et al. 1977; Henschel & Seely 1997). Additionally, species with poor life-history information, such as rare species with few specimens, may lead to inaccurate measurements of intraspecific variation. Allometric scaling laws can be used to overcome these problems. These laws refer to how traits, which can be morphological, physiological or chemical, co-vary with an organism’s body size, often with important ecological and evolutionary implications (Gould 1966). When these scaling laws are utilised to estimate body size or a hypothesised allometric characteristic indirectly using a co-varying morphological trait, therefore circumventing the use of destructive and/or time-consuming methods, we define this as ‘predictive allometry’.

Predictive allometry has emerged across many biological disciplines. The most commonly used co-varying trait used to predict body size is body length, which has been used extensively in fish (e.g. Karachle & Stergiou 2012), mammals (e.g. Trites & Pauly 1998) and both aquatic (e.g. Burgherr & Meyer 1997) and terrestrial invertebrates (e.g. Rogers et al. 1977; Sabo et al. 2002). These models often show considerable predictive power (*R*^*2*^ > 0.9), which has led to the proliferation of multiple models for a wide range of taxa (e.g. there are 26 body length – body size models for Diptera – See Supporting Information). However, when compared, these models show considerably different allometric scaling coefficients both within- and between insect orders (Schoener 1980; Sample et al. 1993; Ganihar 1997; Brady & Noske 2006). Previously, these differences have been attributed to biogeographic factors, such as latitude (Martin et al. 2014) and/or methodological influences such as sampling biases (e.g. the range of sampled body sizes, Sage 1982). Importantly, they have also notably failed to incorporate sexual size dimorphism which is common in invertebrates (Shreeves & Field 2008).

The allometry of functional traits have been shown to influence plant-pollinator interactions, specifically in bees. For example, smaller body size can be associated with preferential activity periods related to available light (Streinzer et al. 2016), whereas larger body size is associated with greater pollen load capacity (e.g. within *Melipona quadrifasciata* colonies, see Ramalho et al. 1998) as well as greater interspecific foraging distances (e.g. Greenleaf et al. 2007). Importantly, body size can both influence and constrain plant-pollinator interactions and trait matching both within and between pollinator groups (Stang et al. 2009; Bartomeus et al. 2016). Therefore, allometric traits central to pollination-related ecological processes both appear and interact at the intra- and interspecific levels. Despite their ubiquity, few predictive models for body size exist for pollinating insects below the ordinal level, with one notable exception. Cane (1987) pioneered a predictive model for bee body size as a function of the intertegular distance (ITD) (the distance between the wing-attachment points on either side of the thorax (See Fig. S1). Importantly, Cane’s allometric model identified the ITD as an important body size proxy and has since been used to establish other ecologically important allometric relationships, primarily at the interspecific level (e.g. foraging distances and bee proboscis length; Greenleaf et al. 2007; Cariveau et al. 2016).

The robustness of the ITD as a body size predictor has not been properly tested. First, the original model is based solely on 20 North American solitary bee species, despite evidence suggesting allometric coefficients can differ significantly between biogeographical regions (Martin et al. 2014). Second, the power of predictive allometric equations in predicting intraspecific variation has not been assessed. Third, sexual size dimorphism is present in 80% of Aculeata (Shreeves & Field 2008), highlighting the need to include sex-specific co-variation. Fourth, body size variation has been repeatedly linked to phylogeny, compelling allometric studies to incorporate species’ evolutionary histories (Garland & Ives 2000; Blomberg et al. 2003). Lastly, other key pollinating taxa, such as hoverflies (Diptera: Syrphidae) lack predictive models that could be used to examine allometric patterns.

These knowledge gaps are largely due to the lack of: (a) a general repository to house and connect all relevant predictive allometric models; (b) large high resolution datasets to build more accurate models that can incorporate co-variates and (c) the absence of an iterative framework, such as those utilised in ecological forecasting (e.g. Dietze et al. 2018; Harris et al. 2018) to continuously update existing models with new datasets, methodologies and technologies. Addressing these key deficiencies will increase model accuracy and applicability of predictive allometry for pollinating insects.

Here, we catalogue pre-existing models for key pollinating insect taxa (Diptera, Hymenoptera and Lepidoptera) and develop new predictive allometric models within an iterative framework for bees and hoverflies that incorporate species evolutionary histories, intraspecific variation and biogeography. These form the basis of a new *R* package, entitled “*pollimetry*”. Specifically, we address the following research questions:

i. Is ITD a robust predictor of inter-specific body size variation for two dominant pollinator taxa, bees and hoverflies?
ii. Does incorporating sexual dimorphism and phylogenetic/taxonomic relatedness when constrained by biogeographic region improve interspecific predictions of pollinator body size by ITD?
iii. Is ITD reliable in predicting intraspecific variation in both bees and hoverflies and what sample size is required to accurately estimate intraspecific body size and co-varying trait values?

## Materials and Methods

### Specimen collection and measurements

We obtained specimens collected in recent field research projects on insect pollinator diversity. We included studies across four continents. In Australia, collections were made in New South Wales, Victoria, Queensland, South Australia and the Northern Territory. In Europe, we amassed specimens from Belgium, UK, Germany, Ireland, Spain and Switzerland. In the Americas, we included collections from Minnesota, USA and Ceará, Brazil. In addition, Cane’s (1987) original data from Alabama, USA was obtained using Engauge Digitizer version 10.6 (Mitchell et al. 2018).

The majority of specimens were dehydrated and weighed within three to six months of collection, although some, in particular, those from Victoria, Australia, Belgium, Switzerland and Cane’s original samples were of variable ages: ranging from one to five years since collection. We excluded damaged specimens. For every specimen, we obtained sample location (latitude and longitude) and taxonomic identity. Full information about specimen identification, deposition locations and used taxonomic resources are provided in the Supporting Information.

### Body size and intertegular distance

Body size was measured as the dry weight in milligrams of each specimen. We therefore refer to body size as dry weight herein for continuity. Dry weight was measured by first dehydrating specimens at 70 °C for at least 24hrs prior to weighing to remove residual humidity and then weighed on an analytical balance to an accuracy of 0.001g. All North American bees as well as small-bodied Australian bees were dehydrated and weighed prior to pinning. For all other specimens, pins were not removed prior to weighing. Instead, we identified the pin type and weighed a sample of 10 – 50 pins per type. The mean weight was then subtracted off the total weight. Pin weight variance was minimal (range of standard errors: 6.3*10^−4^ to 2mg). Intertegular distance was measured in millimetres using a stereo-microscope, either mounted with a calibrated scale or microscope camera. Body length was measured along the lateral side of each specimen with a calibrated scale or microscope camera for Australian, British, German, Irish and Spanish specimens (see Supporting Information for visual representation of ITD and body length measurements).

### Data analysis: Model structures

All analyses were undertaken in *R* (version 3.5.1) (R Core Team 2018). We first assessed the Pearson’s correlation coefficient between ITD and body length. ITD and body length (BL) were highly correlated in both bees (*ρ* = 0.932), and hoverflies (*ρ* = 0.853). We then compared both ITD and body length independently in predicting body size using ordinary least squares (OLS) regression to select the best body weight predictor. ITD was marginally more predictive than BL in estimating dry weight in bees: ITD *R*^*2*^: 0.896; BL *R*^*2*^: 0.877, and considerably better than BL for hoverflies: ITD *R*^*2*^: 0.854; BL *R*^*2*^: 0.796. Hence, we used ITD in the following analyses.

As traditionally performed, we used log-transformed values in the model formulation because allometric relationships are typically described by a power function (*y* = *ax*^*b*^) which is linearised when log-transformed:

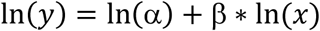

where Y = dry weight, α = intercept, β = allometric co-efficient and *x* = dry weight or body length.

OLS does not allow for the incorporation of random effects or phylogenetic co-variance matrices. Therefore, to incorporate these more complex model structures with the best predictor (i.e. ITD) of dry weight, we specified Bayesian generalised linear mixed models (GLMM) using the *brms* package (version 2.4.0) (Bürkner 2017). Log-transformed dry weight was predicted as a function of the log-transformed ITD in interaction with sex and taxonomic grouping: bee families following Michener (2000) and hoverfly subfamilies following Thompson and Rotheray (1998). We included a nested random effect: species nested within their biogeographic region of origin. A few specimens from five bee species: *Andrena wilkella* (North America), *Halictus rubicundus* (North America), *Lasioglossum leucozonium* (North America), *Anthidium manicatum* (North America) and *Apis mellifera* (Australia), were removed from their introduced ranges (in parentheses) prior to analyses. We call these models taxonomic GLMMs. Both bee and hoverfly models were run for 2000 iterations with a burn-in of 1000. We set Δ to 0.99 and manipulated maximum tree depth between 10 and 20 for individual models to avoid divergent transitions. We fitted each model with weakly informative priors on both fixed and random effects based off our domain expertise; priors are explicitly provided in accompanying R code. Chain convergence was assessed using the 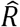 statistic (Gelman & Rubin 1992). Posterior predictive checks were visualised using the *Bayesplot* package (version 1.6.0; Gabry & Mahr 2017).

### Data analysis: Incorporating phylogeny

We explored the influence of phylogenetic relatedness in predicting dry weight for bees only because a well-resolved hoverfly phylogeny was not available. We constructed an applicable phylogeny for our dataset using a bee genera backbone tree (Hedtke et al. 2013). We removed non-represented genera using the *ape* package (version 5.1; Paradis et al. 2004). Species tips were added to genera nodes as simulated pure-birth subtrees using the *phytools* package (version 0.6-44; Revell et al. 2012). This excluded a total of three species (*Flavipanurgus venustus, Protomeliturga turnerea* and *Tetrapedia diversipes*), whose genera weren’t included in Hedtke et al. (2013)’s phylogeny.

As such, we made the explicit assumption that phylogenetic patterns in body size were assessed at and above the genus level. We estimated relative node ages using the mean path lengths method of Britton et al. (2002). We assessed the significance of phylogenetic signal using Pagel’s λ (Pagel 1999) with the *phytools* package (version 0.6-44; Revell et al. 2012). Phylogenetic signal was highly significant for bee ln body size (λ: 0.793, *p* < 0.001) (Fig. 1). Therefore, we implemented a nested phylogenetic generalised linear mixed model (PGLMM) which considered ITD in interaction with intraspecific sexual dimorphism whilst accounting for phylogenetic dependencies through a nested random term: species nested within region (i.e. the nested species term was constrained by the constructed phylogeny). We refer to these models as phylogenetic GLMMs.

**Fig. 1.**
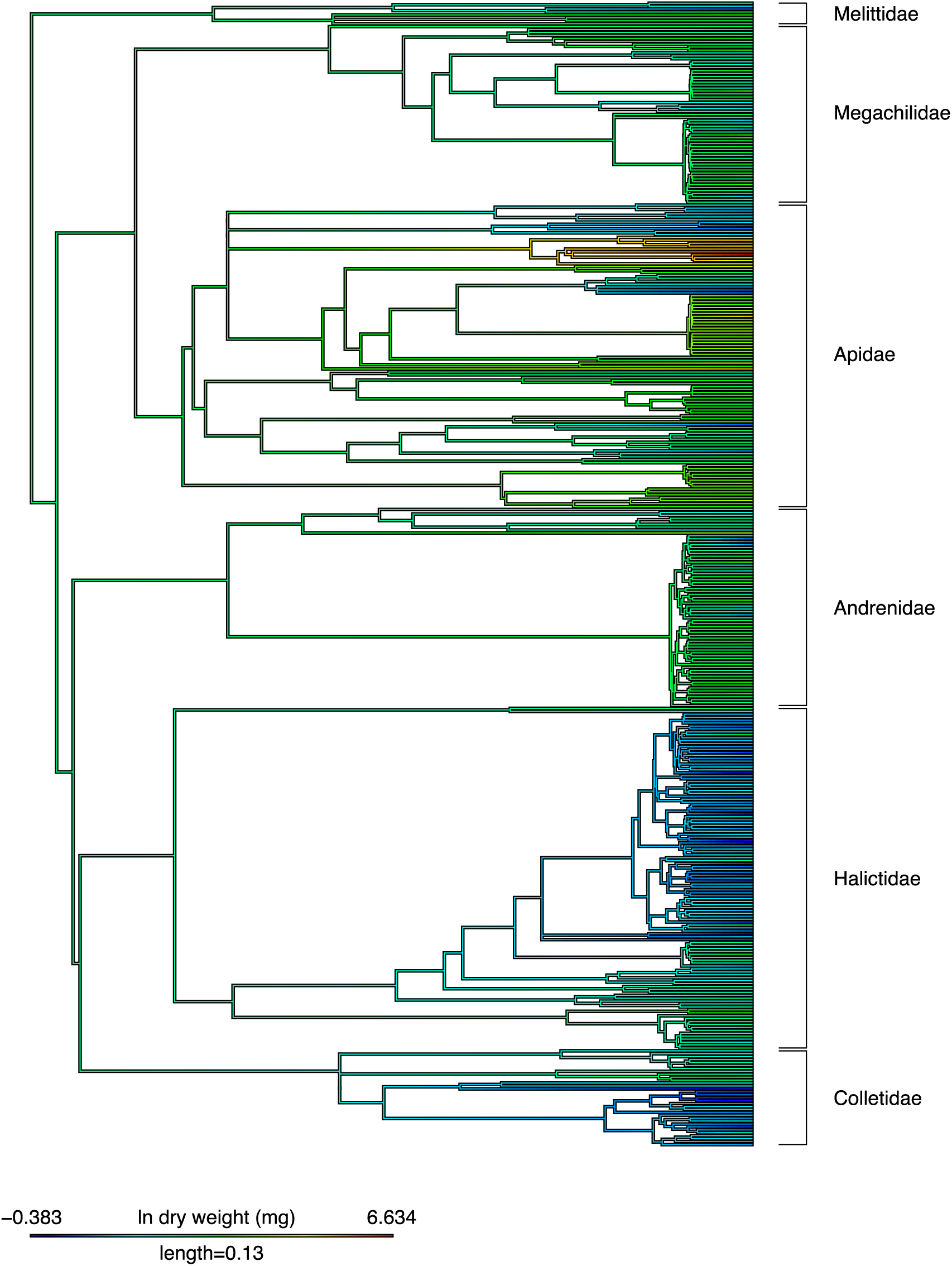
Chronogram of bee genera (Hedtke et al. 2013) with simulated species subtrees. Branch lengths correspond to relative time since divergence. Colour denotes mean ln dry weight (mg) of each bee species.

### Data analysis: Model selection: Bayesian R^2^ and K-fold cross-validation

We first fitted the two full models described above; a taxonomic GLMM and a phylogenetic GLMM. As we were interested in their predictive power, these models were then compared against reduced models (i.e. without sex as either intercepts/slopes) including random effects along with two ITD-only models, one with and one without random terms (Table 1) in order to select the most suitable models for inclusion in the *R* package. We chose to rank our models based upon their Bayesian *R*^*2*^ and K-fold cross-validation (CV) weighting as the Widely-applicable information criterion (WAIC) and Leave-one-out information criterion (LOOIC) were inappropriate due to pWAIC estimates of >0.4 and Pareto k estimates of >0.7 (Gelman et al. 2017; Vehtari et al. 2017). To calculate K-fold CV, species mean datasets were divided into 10 equal sets containing a random subset of species. Each model was then evaluated iteratively upon each k-1 set (training set consisting of nine sets) by comparing the actual and predicted values within the one left out ‘test’ set. This was done repeatedly so each set was both the test set and contained within the training sets from which an information criterion weighting was then calculated.

**Table 1.**
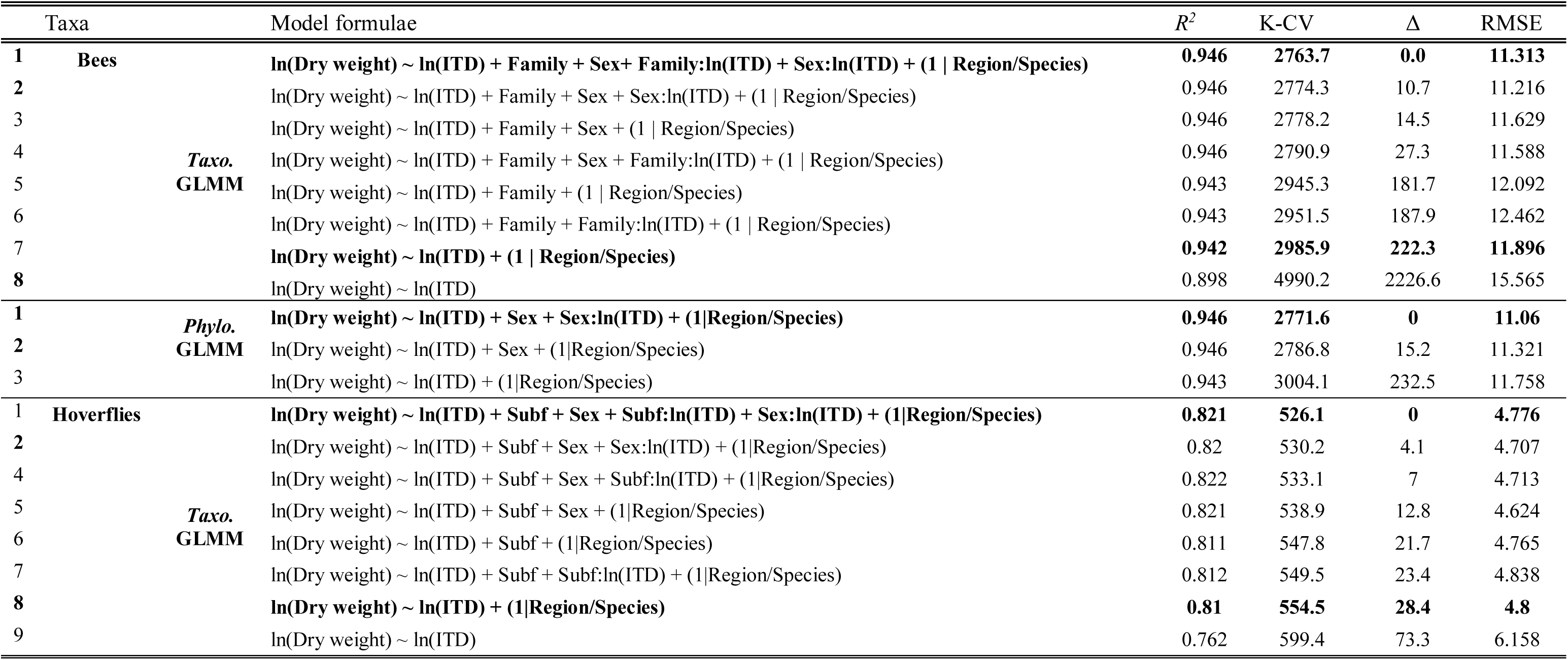
Model selection tables for bee and hoverfly interspecific models. Models in bold are those included in the *R* package. Model types: i) Taxo. GLMM: taxonomic generalised linear mixed models and ii) Phylo GLMM: phylogenetic generalised linear mixed model. lnITD: ln intertegular distance (mm), Subf: Subfamily, *R*^*2*^: Bayesian R^2^, K-CV: K-fold cross validation, Δ: ΔK-fold CV and RMSE: root-mean square error. Estimates of best models are shown in Supporting Information.

### Model comparisons: Root mean square error (RMSE)

We assessed the predictive error of all formulated models on the basis of the root-mean square error (RMSE), which is expressed in the same units of the response variable, between observed-predicted dry weight values and compared these point-estimates of error between our models and predicted values from Cane (1987)’s original model. Lastly, we calculated RMSE for observed-predicted values from existing body length models for both taxa and our body length measurements.

### Data analysis: Intraspecific predictions

We assessed the utility of ITD in predicting intraspecific dry weight variation. For the 10 most abundant bee species of a given sex (nine using females, one using males) and five most abundant hoverfly species (all using females) we tested the utility of ITD in predicting intraspecific body size variation using species-level OLS regression.

To estimate the adequate sample size needed for robust mean trait measures for each bee species, we plotted trait means independently against increasing sample size. We then inferred the adequate sample size whereby variance stabilised within the 95% confidence intervals of the actual sample size.

## Results

### Pre-existing models

We collated 26 predictive allometric models for Diptera, 38 for Hymenoptera and 21 for Lepidoptera groups. We also gathered nine equations for bee foraging distance from two sources (van Nieuwstadt & Iraheta 1996; Greenleaf et al. 2007) and one allometric model for estimating bee tongue length (Cariveau et al. 2016) (See Supporting Information).

### Species and specimen distribution

In total, we measured 391 bee species (4035 specimens) from Australia, Europe, North America and South America and measured 103 hoverfly species (399 specimens) from Australia and Europe (Supporting Information). Six out of seven bee families (all except Stenotritidae) and two hoverfly subfamilies (Syrphinae and Eristalinae) were represented. The mean specimen number per bee species was nine (♀) and five (♂) and ranged from one – 201. In hoverflies, the mean specimen number per species was three for both sexes and ranged from one – 50. In bees, when dry weight variation was visualised across the phylogeny (Fig. 1), large dry weight was most evident within the Apidae, the largest bee in our dataset being the South American *Xylocopa frontalis* (♀ mean weight: 760.75mg). In contrast, Halictid (i.e. *Halictus, Homalictus* and *Lasioglossum* species) and Colletid bees, in particular, the Australian *Euhesma* sp. (♀ mean weight: 0.71mg, ♂ mean weight: 0.66mg) and the European *Hylaeus communis* (♀ mean weight: 6.15mg, ♂ mean weight: 2.76mg) were considerably small.

### Interspecific model selection and performance

All three tested co-variables exhibited significant influences on the allometric scaling of ITD (Fig. 2, Table 1). For bees, both GLMM and PGLMM analyses indicated that models including family or phylogeny and sex in interaction or in addition with ITD, along with our nested random term better predicted dry weight relative to the baseline model (ITD-only model without random term) on the basis of K-fold CV and Bayesian *R*^*2*^(Table 2; Δ*R*^*2*^: 0.046, ΔK-fold CV: 2226.6). However, differences in K-fold CV and Bayesian *R*^*2*^ between the best-fitting taxonomic and phylogenetic models were minimal (Δ*R*^*2*^ <0.001, ΔK-fold CV: 7.92). In hoverflies, incorporating taxonomy and/ sex increased body size predictions relative to the baseline ITD-only models considerably (Δ*R*^*2*^: 0.058, ΔK-fold CV: 73.3).

**Table 2.** Model parameters of intraspecific ln (dry weight) - ln intertegular distance (ITD) relationships. F: F-statistic and degrees of freedom for each model. α: intercept, β: ITD co-efficients ± standard error, *R*^*2*^: Adjusted *R*^*2*^ and P: p-value.

| Taxa | Region | Taxonomic ranking | Species | F<sub>(df)</sub> | $\alpha$ | $\beta$ | $R^2$ | P |
| --- | --- | --- | --- | --- | --- | --- | --- | --- |
| Bee | Europe | Andrenidae: Andreninae | ♀ *Andrena flavipes* | 17.63<sub>(1,70)</sub> | 1.575 ± 0.367 | 1.73 ± 0.412 | 0.189 | <0.001 |
| | Europe | Andrenidae: Andreninae | ♀ *Andrena nigroaenea* | 30.17<sub>(1,50)</sub> | 0.893 ± 0.488 | 2.459 ± 0.448 | 0.364 | <0.001 |
| | North America | Apidae: Apinae | ♂ *Bombus impatiens* | 20.14<sub>(1,66)</sub> | 2.128 ± 0.365 | 1.275 ± 0.284 | 0.222 | <0.001 |
| | Europe | Apidae: Apinae | ♀ *Bombus lapidarius* | 110.2<sub>(1,54)</sub> | 0.277 ± 0.343 | 2.761 ± 0.263 | 0.665 | <0.001 |
| | Europe | Apidae: Apinae | ♀ *Bombus terrestris* | 137.8<sub>(1,81)</sub> | 1.242 ± 0.274 | 2.136 ± 0.182 | 0.625 | <0.001 |
| | Australia | Halictidae: Halictinae | ♀ *Homalictus urbanus* | 6.055<sub>(1,209)</sub> | -0.164 ± 0.033 | 1.166 ± 0.474 | 0.024 | 0.014 |
| | Europe | Halictidae: Halictinae | ♀ *Lasioglossum glabriusculum* | 6.444<sub>(1,47)</sub> | 0.302 ± 0.127 | 2.802 ± 1.104 | 0.102 | 0.014 |
| | Europe | Halictidae: Halictinae | ♀ *Lasioglossum lanarium* | 53.87<sub>(1,61)</sub> | 0.702 ± 0.198 | 2.13 ± 0.29 | 0.46 | <0.001 |
| | Europe | Halictidae: Halictinae | ♀ *Lasioglossum pauxillum* | 37.46<sub>(1,129)</sub> | 0.488 ± 0.057 | 2.715 ± 0.444 | 0.219 | <0.001 |
| | South America | Apidae: Apinae | ♀ *Trigona spinipes* | 0.285<sub>(1,48)</sub> | 2.144 ± 0.243 | 0.287 ± 0.537 | -0.02 | 0.596 |
| Hoverfly | Australia | Syrphidae: Syrphinae | ♀ *Austrosyrphus* sp. 1 | 12.7<sub>(1,30)</sub> | 0.087 ± 0.458 | 2.032 ± 0.57 | 0.274 | 0.001 |
| | Europe | Syrphidae: Syrphinae | ♀ *Episyrphus balteatus* | 0.08<sub>(1,8)</sub> | 1.334 ± 1.885 | 0.885 ± 2.229 | -0.11 | >0.1. |
| | Europe | Syrphidae: Eristalinae | ♀ *Helophilus parallelus* | 14.84<sub>(1,17)</sub> | 0.286 ± 0.857 | 2.485 ± 0.645 | 0.435 | 0.001 |
| | Europe | Syrphidae: Syrphinae | ♀ *Melanostoma scalare* | 6.38<sub>(1,7)</sub> | -2.172 ± 1.324 | 7.619 ± 3.016 | 0.4 | 0.03 |
| | Australia | Syrphidae: Syrphinae | ♀ *Sphaerophoria macrogaster* | 0.04<sub>(1,8)</sub> | 0.361 ± 0.274 | 0.195 ± 0.907 | -0.11 | >0.1. |

**Fig. 2.**
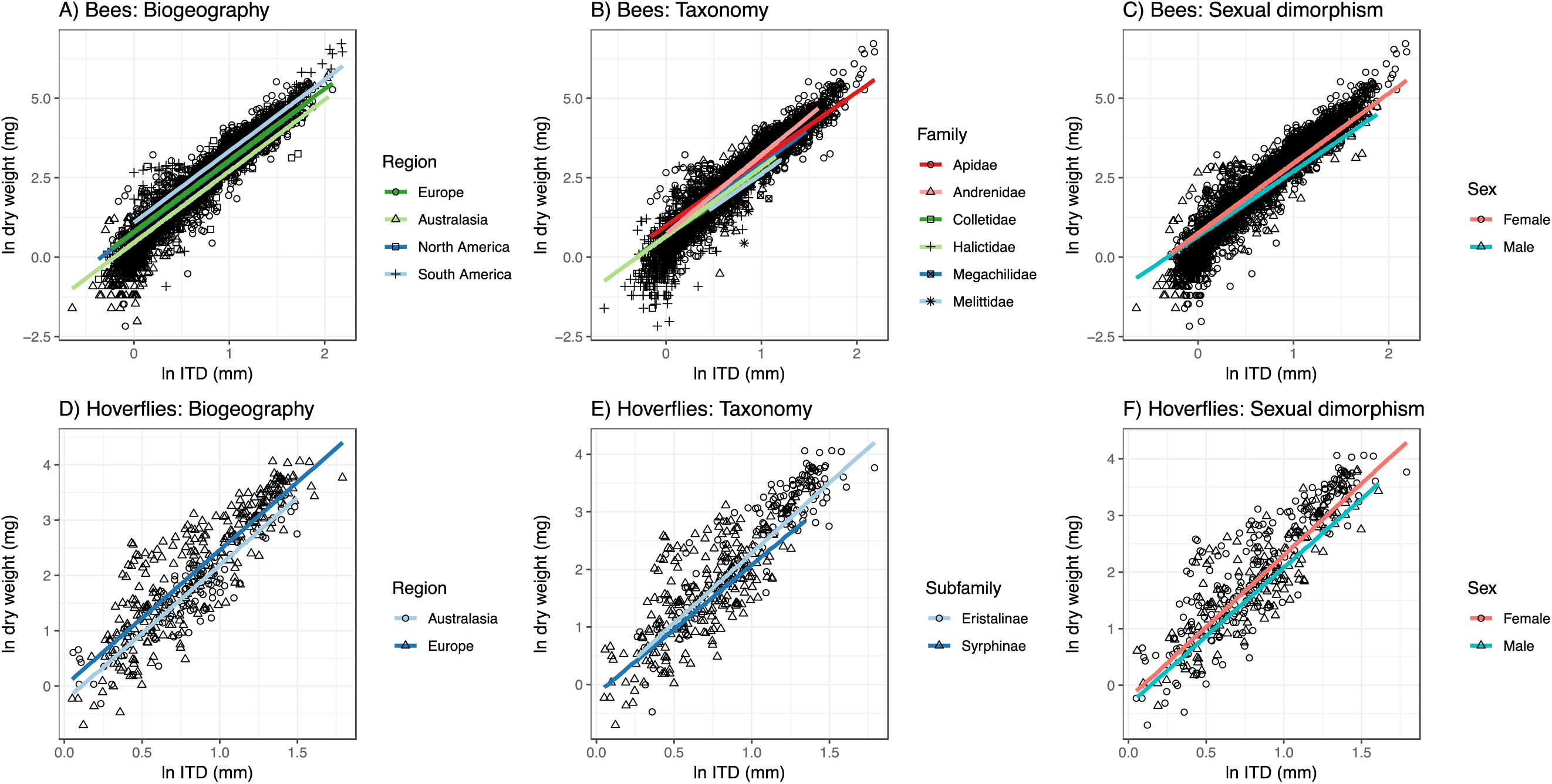
Dry weight (mg) ∼ Intertegular distance (ITD) interspecific relationships. From left to right: influence of biogeographic region, taxonomic grouping and sexual dimorphism. Lines represent the posterior fits from Bayesian generalised linear mixed models. 95% credible intervals are omitted for clarity. See Supporting Information for model co-efficients.

Increases in model performance as a result of incorporating co-variates were most pronounced in bees in terms of root mean square error (RMSE) (Fig. 3). All formulated models outperformed ITD-only models in their predictive precision. RMSE ranged between 10.804 – 12.462mg for both taxonomic and phylogenetic GLMMs. The RMSE for the baseline ITD-only model was 15.565mg, which was near-identical the RMSE for Cane’s (1987) original model: 15.553mg. The RMSE for taxonomic GLMMs for hoverflies ranged from 4.619mg to 4.849mg and all were slightly lower than the RMSE of the baseline ITD-only model (6.179mg). The range of prediction error for ITD was also considerably lower than any pre-existing and applicable model using body length: 36.36mg ± 8.29 for bees and 7.99mg ± 0.69 for hoverflies.

**Fig. 3.**
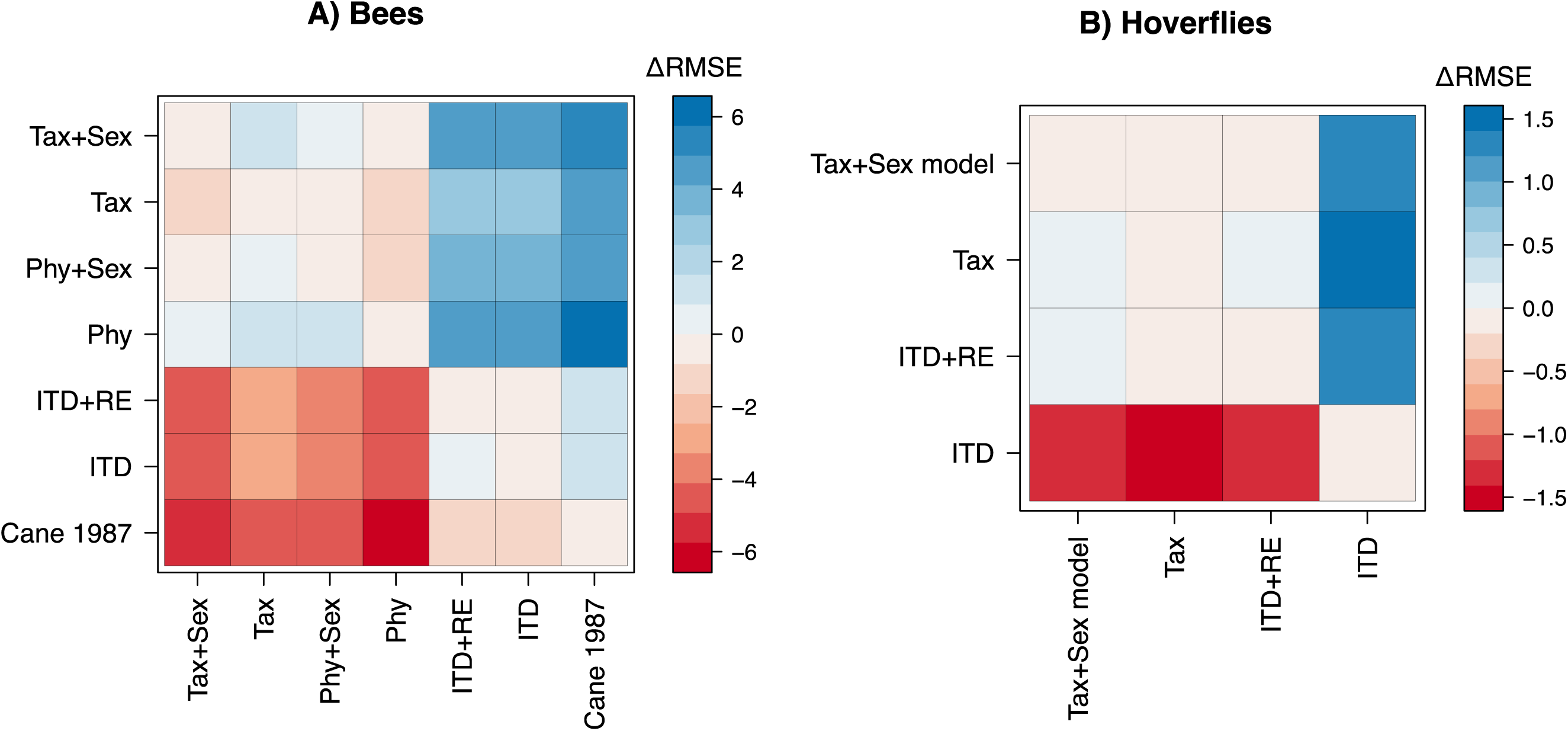
Pairwise comparisons of Δ root mean square error (RMSE) in milligrams between bee and hoverfly models. Negative values denote that models on x axis have lower precision, whereas positive values signify higher precision. Tax+Sex: Full taxonomic model, Tax: Reduced taxonomic model, Phy+Sex: Full phylogenetic model, Phy: Reduced phylogenetic model, ITD+RE: ITD mixed effect model, ITD: ITD fixed effect model. Cane (1987)’s original model for bees.

**Fig. 4.**
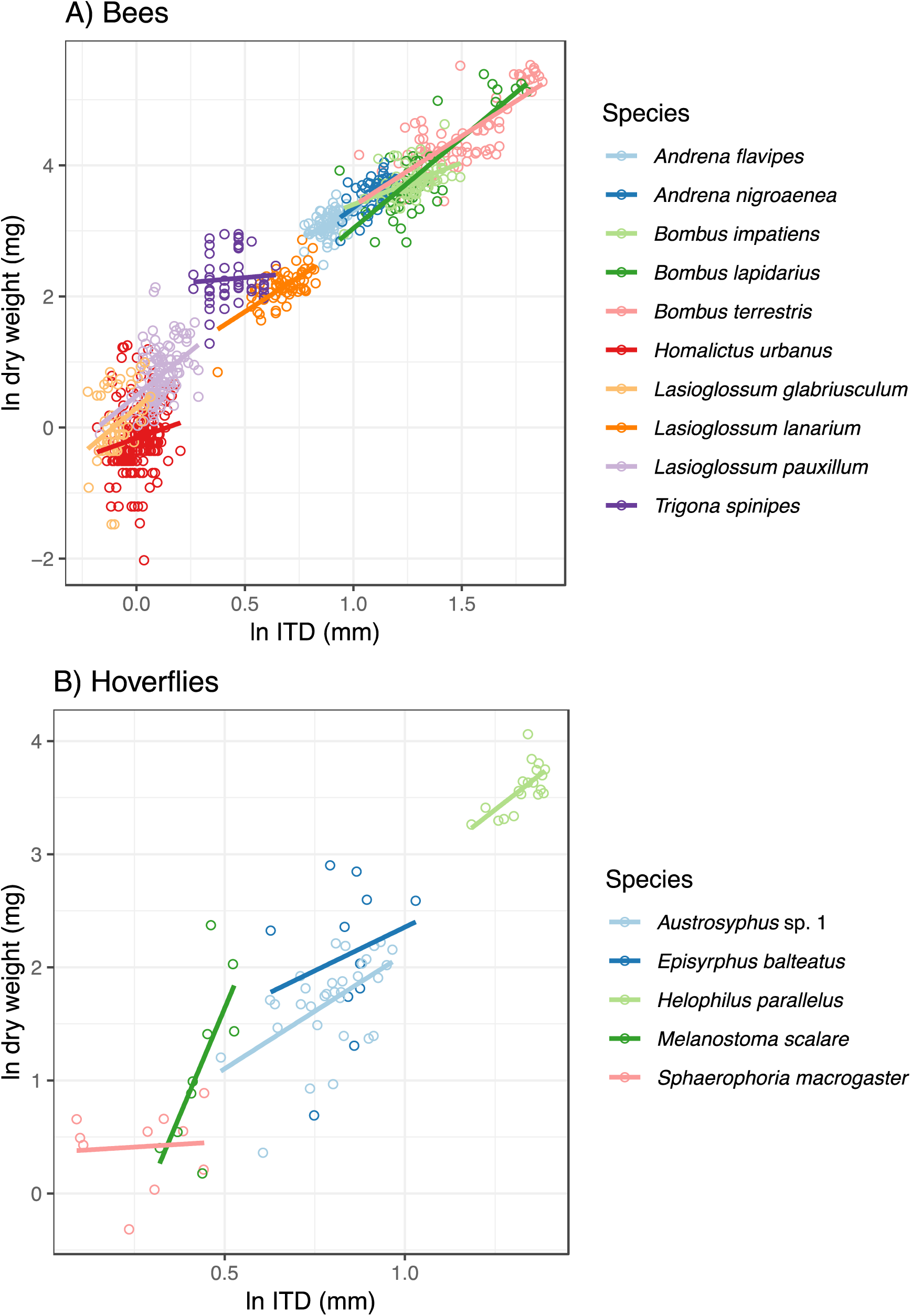
Intraspecific predictions of female* dry weight with intertegular distance (ITD)). Lines denote line of best fit from OLS regression. *Except for Bombus impatiens.

### Intra-specific predictions

Across the 10 most abundant species of bees (♀ *Andrena flavipes*, ♀ *A. nigroaenea*, ♂ *Bombus impatiens*, ♀ *B. lapidarius*, ♀ *B. terrestris*, ♀ *Homalictus urbanus*, ♀ *Lasioglossum glabriusculum*, ♀ *L. lanarium*, ♀ *L. pauxillum* and ♀ *Trigona spinipes*) and five most abundant hoverflies (♀ *Austrosyrphus* sp. 1, ♀ *Episyrphus balteatus*, ♀ *Helophilus parallelus*, ♀ *Melanostoma scalare* and ♀ *Sphaerophoria macrogaster*), the strength of intraspecific predictions of body size using ITD varied considerably (Table 3; Fig. 3). All bee species exhibited a significant relationship, however the adjusted-*R*^*2*^ differed considerably from 0.02 in *Homalictus urbanus* to 0.66 for *Bombus lapidarius*. Similarly, three of five hoverfly species, *Austrosyrphus* sp., *Helophilus parallelus and Melanostoma scalare* exhibited a significant relationship. In order to accurately determine mean ITD and dry weight values for bees, a sample size of 20-30 specimens is required for trait values to stabilise within the 95% confidence intervals of the total sample size (See Supporting Information).

### Summary of R package functions

The developed *R* package, ‘*pollimetry’*, integrates models for estimating body size (i.e. dry weight) in bees and hoverflies using the ITD and co-variates (see Table 2), which were parameterized with the enclosed dataset, into a wrapper function that returns body size estimates, along with standard error and 95% credible intervals. In addition, *pollimetry* includes functions for estimating pollinator dry weight using pre-existing models which utilise the following co-varying traits: body length, head width and body length * body width; see Supporting Information). The *R* package also includes functions for estimating bee foraging distances using the ITD (Greenleaf et al. 2007) or head width (van Nieuwstadt & Iraheta 1996), as well as models for estimating bee tongue length using the ITD and taxonomic family (Cariveau et al. 2016). The equations will be updated in future package releases as novel data become available and models are re-fit to these new data.

## Discussion

We present the most comprehensive examination of allometric scaling for predictive means for two important pollinating insect taxa: bees and hoverflies. We propose an iterative framework to develop and test this suite of highly predictive dynamic allometric models that consider allometric scaling variation attributable to phylogenetic relatedness, sexual dimorphism and biogeographic differentiation.

Incorporating phylogenetic information is a cornerstone of comparative biological analyses, especially in studies concerning body size variation. Phylogenetic signal in body size variation has been inferred in a number of vertebrate and invertebrate groups (Ashton 2004). Failing to account for dependent phylogenetic patterns is argued to heighten the risk of inaccurate predictions (Martins et al. 2002; Garland et al. 2005). In our study, both PGLMM and GLMM models were comparable in terms of predictive power as well as parameter values. Interestingly, taxonomic and phylogenetic GLMM models were near-identical in all model rankings (Bayesian *R*^*2*^, K-fold CV and RMSE), demonstrating that differential allometric scaling is present at/or below the familial level. These results suggest that predictive inferences of body size above the family level lack accuracy and generalisability.

Where the aim is prediction, GLMMs incorporating taxonomic groupings without considering phylogeny are more practical given well-resolved phylogenies are lacking for most groups (e.g. one can predict allometric relationships for non-represented species). A further advantage of using taxonomic groupings over phylogeny is that they provide easy-to-interpret regression intercepts and/or slopes as opposed to a phylogenetic co-variance matrix. Therefore, for bees, we confirm that incorporating taxonomy is predictively equivalent in predicting allometric scaling relationships where phylogenetic information is unavailable. Importantly, this uniformity between taxonomic and phylogenetic models may not exist for other taxa with either high paraphyly or low correspondence between taxonomy and phylogeny. In hoverflies, including subfamily was less informative, yet still retained, in describing body size variation, potentially due to their lower taxonomic ranking. In essence, our results suggest that where previous studies have used taxonomy (i.e. bee families in Cariveau et al. 2016), results are predictively comparable to incorporating phylogeny.

Sex was retained as an integral predictor either in addition or in interaction with ITD for both taxa. Sexual size dimorphism (SSD) is common among insects. In both Diptera and Hymenoptera, 80% of previously-studied species exhibit female-biased SSD including in Apoidea and Syrphidae (Shreeves & Field 2008; Francuski et al. 2011). Female-biased SSD is hypothesised to be a result of greater fitness and increased fecundity as a result of larger female body size (Stillwell et al. 2010). In bees, SSD is attributed to the physical requirements of nest provisioning and construction (Shreeves & Field 2008). This suggests that intraspecific sex differences in the allometric scaling of ITD may reflect the presence of sex-specific morphologies such as the presence of specialised morphological structures for resource collection (i.e. scopal hairs and corbiculae) as well as self-preservation (i.e. a stinger) in female bees.

In hoverflies, SSD was also notably female-biased, with sex retained as an important body size predictor in conjunction with the ITD. However, few examples of morphological sexual dimorphism exist. In both taxa, including sex increased model precision by <4.25-1.38mg RMSE, highlighting the predictive accuracy of the ITD even when sex is not considered. Therefore, failing to incorporate sex in predictions will only introduce a subtle error. Sex is easily identifiable in both bees and hoverflies. Therefore, we recommend its inclusion if predictive allometries are used as many ecologically relevant allometric traits are sex-related (e.g. flight distances; Kraus et al. 2009).

Few previous studies have assessed the utility of predictive models in describing intrageneric or intraspecific allometric traits (e.g. Hagen & Dupont 2013; Cariveau et al. 2016). Our results suggest that intraspecific body size variation is difficult to predict accurately using co-varying traits such as the ITD. In particular, the large variation in predictive power suggests that it is sensitive to environmental conditions and/or sample sizes. Adult body size variation in holometabolous insects is a direct result of diet and environment during ontogeny and larval development (Davidowitz et al. 2004). For example, within *Bombus* species, brood sizes increase throughout the season in response to colony population increases (Inoue 1992). These intra-specific patterns raise the question of how many individuals are necessary to measure to accurately capture species’ mean trait values. Based on our examination of trait-sample size relationships we can provide a recommendation that measuring 20-30 specimens per species will lead to accurate estimation intraspecific body size and morphological trait values. By applying our iterative framework, we aim to reduce the noise in interspecific models due to low sample sizes in some species by incorporating novel data sets.

Terrestrial invertebrates show considerable biogeographic variation in body shape and size. While previous studies have compared predictive allometries between biogeographical regions either independently (Schoener 1980) or within a meta-analytical framework (Martin et al 2014), we chose to represent biogeographical variation within a random effect structure. This makes these models broadly applicable and not biogeographically restricted in utility. Observed biogeographical differences within this study likely arise from differing species diversification patterns as well as from sampling biases, such as variation in commonality among species. Therefore, discerning hypotheses that explain biogeographic variation in the allometric scaling of ITD is problematic. However, it is clear that the influence of biogeography appears alongside species’ evolutionary histories and intraspecific variation.

By incorporating phylogeny or taxonomy, sexual dimorphism and biogeographic random effects we improved model predictions and reduced the limitations of traditional predictive allometry. These three predictors represent fundamentally-related causes of body size variation in pollinating insects. In consideration of the multiple metrics (i.e. Bayesian *R*^*2*^, K-fold CV, and RMSE) used in model selection and performance, we provide multiple, near-equally accurate predictive models. This is important as research questions may not garner investigation of sex-related allometric differences and may occur outside the included biogeographic regions. Therefore, disseminating the most appropriate allometric model becomes a hypothesis-driven formula that should consider and then discount each examined factor. Importantly, given the high resolution across our described models and large sample size of specimens within our study, our models will improve body size predictions relative to pre-existing models even when considering only ITD. After accounting for biogeographical and species-level effects, failing to incorporate sex or phylogeny/taxonomy will not result in considerable error (see Fig. 3) although we endorse their use as it enables more meaningful analyses. Lastly, we caution the use of ordinal-level predictive models as allometric constraints are ubiquitous at the familial level (See Fig. 1).

### Conclusions and implications

The accompanying *R* package, “*pollimetry*”, provides a user-friendly interface to estimate pollinator body size (as dry weight) and modelled allometric traits. Practical predictive allometric libraries require multiple models that are continually updated when novel datasets become available. This will enable robust investigation of other allometric traits at both intra- or inter-specific levels. The consequences of body size variation are ubiquitous within pollination research, yet few have utilised allometric theory in studying pollinating taxa beyond bees. Adding hoverflies is an important first step, yet this comprehensive approach to predictive allometric model development should be applied to other pollinating taxa, such as moths and butterflies. The iterative framework used herein heralds a dynamic new direction for predictive allometry and will provide more accurate predictions through hypothesis-led model choice, testing and investigation in allometric research.

## Acknowledgements

The authors would like to thank K. Freidrich, A. Irber, L. Kirkland, J. Krauss, L. Kuehn, J. B. Lanuza and J. Lumbers for providing specimens, R. Bärfuss for identifying Swiss hoverflies, S. Gerber, M. Herrmann and A. Müller for identifying Swiss bees, K. Mandery for identifying German specimens, S. Wright for identifying Australian hoverflies, O. Aguado for identifying Spanish specimens and A. Pauly for identifying Belgian Halictidae. Finally, we thank M. Betancourt for statistical advice. This study was funded by the BeeFun project (PCIG14-GA-2013-631653), a CSIRO PhD top-up scholarship and an Ian Potter Foundation PhD scholarship grant to LKK and an Australian Research Council Discovery Early Career Researcher Award DE170101349 to RR.

## Data availability

All data including *R* code and the *R* package are available here: https://github.com/liamkendall/pollimetryDOI:10.5281/zenodo.1313905

## Author contributions

IB, LKK, VG and RR conceived the study. LKK, VG, JR and MH collected Australian specimens. LKK measured Australian, German and Swiss specimens. LKK and MH identified Australian bees. ZMP identified North American specimens. LR collected, identified and measured Irish specimens. JMM collected and identified British specimens. FPM collected, identified and measured Spanish specimens. NJV and SPMR collected, identified and measured Belgian specimens. MA and LS collected and identified Swiss specimens. LKK, IB, and VG devised and undertook all data analyses. LKK and IB formulated and wrote the R package. LKK wrote the manuscript and all authors contributed significantly to the final manuscript.

## Supporting Information

### Description of pre-existing models

In addition to developing new predictive allometric models for bees and hoverflies, we selected the three key pollinating insect orders: Diptera, Hymenoptera and Lepidoptera and collated all known predictive allometric models for those orders, regardless of whether they are acknowledged pollinators. Lepidoptera were not included in primary within-text analyses for logistic reasons and low abundances across sourced research projects. From an initial literature search, we obtained the publications analysed by Martin et al. (2014). We then reviewed each publication individually, including their references and citations for additional models.

Diptera: 26 predictive allometric models for Diptera were collated (Table S1A). Eleven models were reported for the entire order, including nine without any taxonomic breakdown of samples used. Twelve models were collated for the three main suborders Nematocera (6), Brachycera (4) and Cycllorapha (2) and two for specific families; Asilidae and Bombyliidae.

Hymenoptera: 38 predictive allometric models for Hymenoptera were collated (Table S1B). These included eight models for the entire order, ten for Formicidae and seven for all Hymenoptera excluding Formicidae. There are three models for Vespidae and two models for Apidae (Cane 1987 & Sabo et al. (2002). Sample et al’s (1993) body length and body length * body width models are provided for Braconidae, Ichneumonidae, Halictidae and Pompilidae.

Lepidoptera: 21 predictive allometric models for Lepidoptera were collated (Table S1C). This includes 13 with varying taxa and without lower classifications. Hodar (1997) provides specific models for Heterocera (moths) and Ropalocera (butterflies). Sample et al. (1993) provide body length and body length * body width models for Microlepidoptera and two moth families: Geometridae and Arctiidae.

**Table S1A.** Allometric models for Diptera. Measure denotes trait measurement (BL = Body length, BW = Body width). Reg = regression type(L = Linear regression. MA = Major axis regression or OLS = Ordinary Least Squares regression). Type denotes slope (EXP = exponential model, PF = power function). Models are present in the form of y = ln(*α*) + ln(*β*) * *x* unless Type noted with *. ** = Included body width as well as length.

| Source | Tax. grouping | Sample size | Biog. region | Measure | Range (mm) | Reg. | Type | Model |  |  |  |
| --- | --- | --- | --- | --- | --- | --- | --- | --- | --- | --- | --- |
| | | (Families: species) | | | | | | $\alpha \pm \text{S.E.}$ | $B \pm \text{S.E.}$ | Resi. SE | $R^2$ |
| Rogers et al. (1977) | NA | (NA:84) | USA | BL | 0.9-34 | OLS | PF | $-3.298 \pm 0.115$ | $2.366 \pm 0.078$ | 0.57 | **0.96** |
| Schoener (1980) | NA | (NA:107) | Costa Rica | BL | NA | OLS | PF | $-2.603 \pm 0.0688$ | $1.64 \pm 0.1224$ | NA | 0.80 |
| " | NA | (NA:124) | " | " | " | " | " | $-2.688 \pm 0.051$ | $1.59 \pm 0.1173$ | NA | 0.78 |
| " | NA | (NA:171) | USA | " | " | " | " | $-3.816 \pm 0.561$ | $2.42 \pm 0.0969$ | NA | 0.89 |
| Gowing and Recher (1984) | NA | (NA:100) | Australia | BL | 2-11 | OLS | PF | $3.653 \pm 0.129$ | $2.546 \pm 0.071$ | 0.37 | **0.93** |
| Sample et al. (1993) | Combined | (15:257) | USA | BL | 2.9-23.65 | OLS | PF | $-3.184 \pm 0.184$ | $2.213 \pm 0.085$ | NA | 0.85 |
| " | " | " | " | BL*BW | " | " | " | $-2.197 \pm 0.089$ | $1.309 \pm 0.03$ | " | **0.94** |
| " | BIB, SCI, TIP | (3:46) | " | BL | 3.55-23.65 | " | " | $-3.675 \pm 0.23$ | $2.212 \pm 0.141$ | " | **0.92** |
| " | " | " | " | BL*BW | " | " | " | $-2.217 \pm 0.205$ | $1.288 \pm 0.071$ | " | **0.94** |
| " | ASI, DOL, EMP, RHA, STR, THE | (6:80) | " | BL | 2.9-17.99 | " | " | $-3.374 \pm 0.230$ | $2.158 \pm 0.101$ | " | **0.92** |
| " | " | " | " | BL*BW | " | " | " | $-2.2 \pm 0.147$ | $1.259 \pm 0.049$ | " | **0.95** |
| " | CAL, LAU, MUS, OTI, SYR, TAC | (6:119) | " | BL | 2.9-15.65 | " | " | $-3.619 \pm 0.212$ | $2.632 \pm 0.101$ | " | **0.92** |
| " | " | " | " | BL*BW | " | " | " | $-2.02 \pm 0.131$ | $1.298 \pm 0.042$ | " | **0.94** |
| Hodar (1997) | BRA | (NA:26) | Spain | HW | NA | OLS | PF | $0.655 \pm 0.105$ | $2.526 \pm 0.139$ | 0.47 | **0.93** |
| " | NEM | (NA:10) | " | " | NA | " | " | $3.942 \pm 0.259$ | $3.106 \pm 0.278$ | 0.55 | **0.94** |
| Ganihar (1997) | NA | (NA:20) | India | BL | NA | OLS | PF | $-3.4294 \pm 0.01994$ | $2.5943 \pm 0.0334$ | 0.03 | **0.99** |
| Johnson and Strong (2000) | NA | (NA:75) | Jamaica | BL | 1-12.5 | OLS | PF | $-2.462 \pm 0.196$ | $1.881 \pm 0.146$ | NA | 0.83 |
| " | NEM | (NA:21) | " | " | 1-4.8 | " | " | $-2.562 \pm 0.244$ | $1.373 \pm 0.207$ | " | 0.84 |
| " | NEM exc. | (NA:54) | " | " | 1.2-12.5 | " | " | $-2.105 \pm 0.178$ | $1.805 \pm 0.124$ | " | 0.89 |
| Sabo et al. (2002) | BRA | NA | USA | BL | NA | OLS | PF* | $0.006 \pm 0.007$ | $3.05 \pm 0.36$ | NA | 0.85 |
| " | NEM | NA | " | " | " | " | " | $0.1 \pm 0.06$ | $1.57 \pm 0.2$ | " | **0.90** |
| " | ASI | (1:9) | " | " | " | " | " | $0.38 \pm 2.625$ | $1.5 \pm 2.469$ | " | 0.74 |
| " | BOM | (1:10) | " | " | " | " | " | $0.007 \pm 0.011$ | $3.337 \pm 0.676$ | " | **0.95** |
| Brady and Noske (2006) | NA | (NA:9) | Australia | BL | 2-28 | OLS | L* | $-0.041 \pm 0.004$ | $0.010 \pm 0.001$ | 0.02 | 0.84 |
| Wardhaugh (2013) | NA | (NA:16) | Australia | BL | NA | MA | PF | $-3.29 \pm 0.45$ | $2.65 \pm 0.36$ | NA | 0.72 |
| " | " | " | " | BL * BW | NA | " | " | $-1.91 \pm 0.19$ | $1.22 \pm 0.11$ | " | 0.87 |

BIB =Bibionidae, SCI = Sciaridae, TIP = Tipulidae, ASI= Asilidae, BOM: Bombyliidae, DOL = Dolichopodidae, EMP = Empididae, RHA = Rhagionidae, STR = Stratiomyidae, THE = Therevidae, CAL = Calliphoridae, LAU = Lauxaniidae, MUS = Muscidae, OTI = Otitidae, SYR = Syrphidae, TAC = Tachinidae. NEM = Nematocera, BRA= Brachycera, CYC = Cyclorrapha

**Table S1B.**
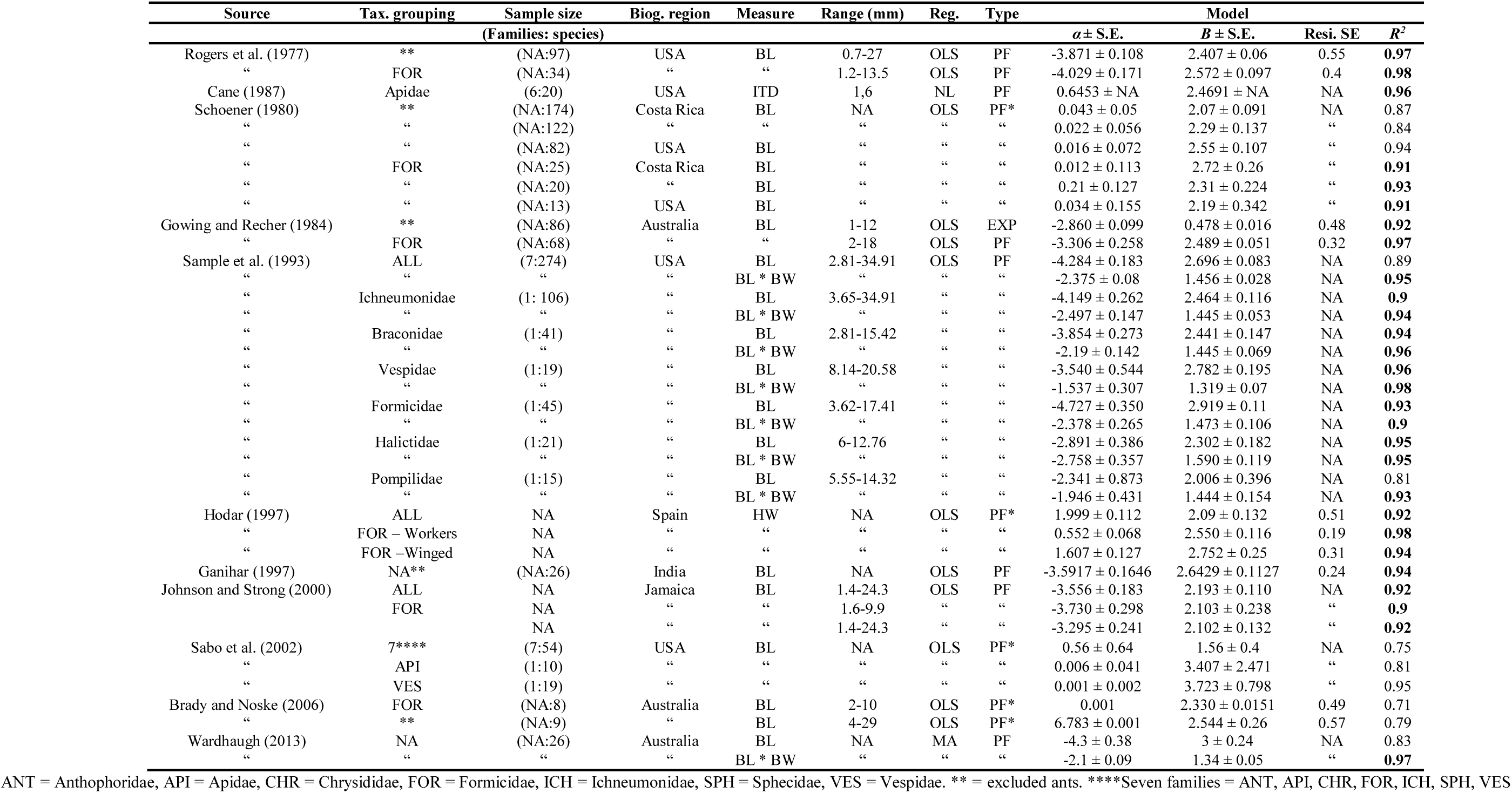
Allometric models for Hymenoptera. Measure denotes trait measurement (BL = Body length, BW = Body width, ITD = Intertegular distance). Reg = regression type (L = Linear regression. MA = Major axis regression or OLS = Ordinary Least Squares regression). Type denotes slope (EXP = exponential model, PF = power function). Models are present in the form of y = ln(*α*) + *β \** In(*x*) unless Type noted with *. ** = Included body width as well as length.

**Table S1C.** Allometric models for Lepidoptera. Measure denotes trait measurement (BL = Body length, BW = Body width). Reg = regression type (MA = Major axis regression, OLS = Ordinary Least Squares regression). Type denotes slope (EXP = exponential model, PF = power function). Models are present in the form of *y* = ln (*α*) + ln (*β*) * *x* unless Type noted with *. ** = Included body width as well as length.

| Source | Tax. grouping | Sample size | Biog. region | Measure | Range (mm) | Reg. | Type | Model |  |  |  |
| --- | --- | --- | --- | --- | --- | --- | --- | --- | --- | --- | --- |
| | | (Families: species) | | | | | | $\alpha \pm \text{S.E.}$ | $\beta \pm \text{S.E.}$ | Resi. SE | $R^2$ |
| Rogers et al. (1977) | NA | (NA:22) | USA | BL | 1.6-17 | OLS | PF | $-4.037 \pm 0.133$ | $2.903 \pm 0.08$ | 0.31 | **0.99** |
| Sample et al. (1993) | ALL | (NA:384) | USA | BL | 2.76-40.73 | OLS | PF | $-5.036 \pm 0.157$ | $3.122 \pm 0.064$ | NA | **0.93** |
| " | " | " | " | BL * BW | " | " | " | $-2.607 \pm 0.088$ | $1.457 \pm 0.024$ | " | **0.95** |
| " | MIC. | (NA:46) | " | BL | 2.76-10.6 | " | " | $-4.913 \pm 0.325$ | $2.918 \pm 0.169$ | " | **0.93** |
| " | " | " | " | BL * BW | " | " | " | $-2.715 \pm 0.199$ | $1.395 \pm 0.08$ | " | **0.93** |
| " | GEO | (1:58) | " | BL | 6.45-21.70 | " | " | $-4.172 \pm 0.411$ | $2.628 \pm 0.167$ | " | **0.9** |
| " | " | " | " | BL * BW | " | " | " | $-2.343 \pm 0.283$ | $1.387 \pm 0.084$ | " | **0.91** |
| " | ARC | (1:60) | " | BL | 5.05-20.06 | " | " | $-3.755 \pm 0.242$ | $2.658 \pm 0.105$ | " | **0.96** |
| " | " | " | " | BL * BW | " | " | " | $-1.658 \pm 0.148$ | $1.222 \pm 0.044$ | " | **0.96** |
| Hodar (1996) | HET | (NA:10) | Spain | HW | NA | OLS | PF | $2.053 \pm 0.25$ | $2.804 \pm 0.236$ | 0.49 | 0.95 |
| " | ROP | (NA:10) | " | HW | NA | " | " | $1.634 \pm 0.46$ | $2.793 \pm 0.446$ | 0.48 | 0.83 |
| Ganihar et al. (1997) | NA | (NA:10) | India | BL | NA | OLS | PF | $-4.7915 \pm 0.751$ | $2.8585 \pm 0.257$ | 0.46 | **0.93** |
| Johnson and Strong (2000) | NA | (NA:40) | Jamaica | BL | 2.2-18.6 | OLS | PF | $-3.268 \pm 0.255$ | $2.243 \pm 0.130$ | NA | **0.94** |
| Schoener (1980) | NA | (NA:29) | Costa Rica | BL | NA | OLS | PF* | $0.026 \pm 0.186$ | $2.55 \pm 0.571$ | NA | **0.96** |
| " | " | (NA:7) | " | " | " | " | " | $0.078 \pm 0.139$ | $1.32 \pm 0.683$ | " | 0.75 |
| " | " | (NA:18) | USA | " | " | " | " | $0.014 \pm 0.18673$ | $2.55 \pm 0.571$ | " | 0.77 |
| Brady and Noske (2006) | NA | (NA:6) | Australia | BL | 7.34 | OLS | PF* | 0.001 | $2.313 \pm 0.223$ | 0.4 | 0.81 |
| Wardhaugh (2013) | NA | (NA:11) | Australia | BL | NA | MA | PF | $-3.83 \pm 0.41$ | $2.77 \pm 0.27$ | NA | 0.83 |
| Wardhaugh (2013) | " | " | " | BL * BW | " | " | " | $-2.1 \pm 0.21$ | $1.37 \pm 0.11$ | " | 0.88 |

HET = Heterocera, ROP = Ropalocera, MIC = Microlepidoptera, GEO = Geometridae, ARC = Arctiidae

**Table S2.**
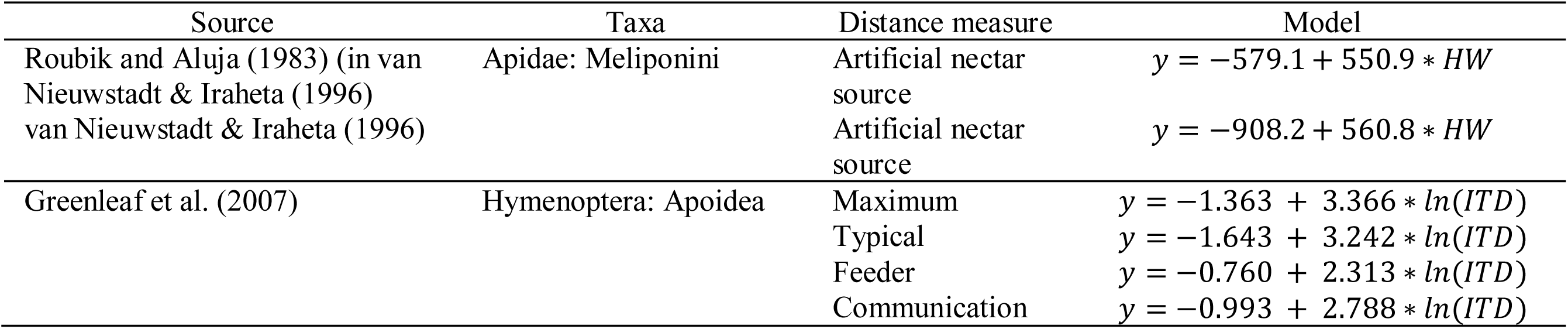
Predictive allometries for bee foraging distance. HW: Head width, IT: Intertegular distance.

**Table S3.**
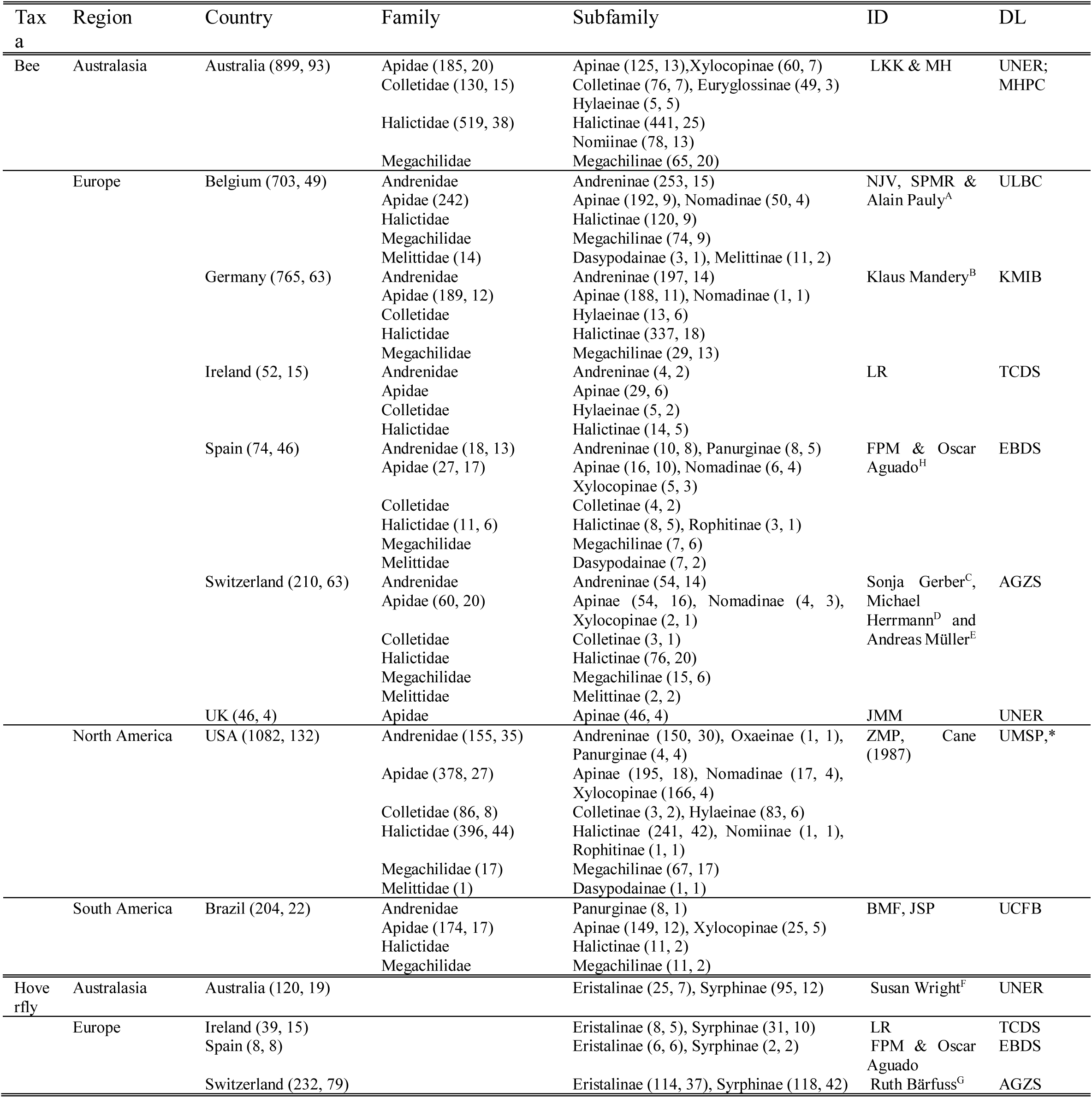
Distribution of included specimens. Numbers in parenthesis denote total specimens and species per country, family and/or subfamily. Exact sampling locations are available in the included dataset. ID: Specimen identifier. Either study author initials or full name and affiliation. DL: Specimen deposition location. Numbers refer to author affiliations or institution address is provided. * All excluding Jim Cane’s specimens (see Cane 1987).

*Taxonomist affiliations*: **A**: Institut royal des Sciences naturelles de Belgique, O.D. Taxonomie & Phylogénie, Rue Vautier 29, 1000 Bruxelles, Belgium. **B**: Institut für Biodiversitätsinformation e.V. Geschwister-Scholl-Str. 6 96106 Ebern, Germany. **C**: Drosera Ecologie Appliquée SA Chemin de la Poudrière 36 1950 Sion Switzerland. **D**: WAB-Mauerbienenzucht Sonnentauweg 47 78467 Konstanz Germany. **E**: Natur Umwelt Wissen GmbH Bergstrasse 162 8032 Zürich Switzerland. **F**: Queensland Museum, PO Box 3300, South Brisbane BC, Queensland 4101, Australia. **G**. Ruth Bärfuss Feldstrasse 7 8625 Gossau ZH Switzerland. **H**: Freelance/no affiliation.

*Specimen deposition locations*: **AGZS**: Agroscope, Agroecology and Environment, Zürich, Switzerland. **EBDS**: Estación Biológica de Doñana Collection, Sevilla, Spain. **KMIB**: Klaus Mandery’s collection, Institut für Biodiversitätsinformation, Bern, Germany. **MHPC**: Mark Hall’s personal collection, Australia. **TCDS**: Stout Lab, Trinity College, Dublin, Ireland. **UCFB**: Bee Laboratory Collection, Federal University of Ceará, Fortaleza, Brazil. **ULBC**: Agroecology Lab reference collection, Université libre de Bruxelles (ULB), Belgium. **UMSP**: University of Minnesota Insect Collection, USA. **UNER**: Rader Lab Insect Collection, University of New England, Armidale, Australia.

---

**Table S4A.** Bees: Model parameters of best-fitting taxonomic GLMM and phylogenetic GLMM. Post.: Posterior mean estimate (95% credible intervals). ESS: Effective sample size. Taxo-GLMM model formula: ln Dry weight ∼ ln ITD + Family + Family:ln(ITD) + Sex + Sex:ln(ITD) + (1|Region/Species). Phylo-GLMM formula: ln Dry weight ∼ ln ITD + Sex + Sex:ln(ITD) + (1|Region/Species).

| Effect type |  | Taxo. GLMM |  |  | Phylo. GLMM |  |
| --- | --- | --- | --- | --- | --- | --- |
|  |  | Parameters | Post. | ESS | Post. | ESS |
| Fixed | Intercept |  | 1.05 (0.64 – 1.37) | 873 | 0.81 (0.32 – 1.25) | 660 |
|  | lnITD |  | 2.09 (2.00 – 2.19) | 893 | 2.03 (1.94 – 2.11) | 1888 |
|  | Sex | Male | -0.1 (-0.15 – -0.05) | 2552 | -0.13 (-0.17 – -0.08) | 3020 |
|  |  | Male:lnITD | -0.15 (-0.21 – -0.09) | 3069 | -0.14 (-0.2 – -0.08) | 3263 |
|  | Family | Andrenidae | -0.08 (-0.27 – 0.10) | 879 | NA | NA |
|  |  | Colletidae | -0.33 (-0.52 – -0.13) | 629 | NA | NA |
|  |  | Halictidae | -0.35 (-0.49 – -0.21) | 632 | NA | NA |
|  |  | Megachilidae | -0.17 (-0.40 – 0.05) | 1024 | NA | NA |
|  |  | Melittidae | -0.47 (-0.97 – 0.03) | 2548 | NA | NA |
|  |  | Andrenidae:lnITD | 0.10 (-0.09 – 0.28) | 1265 | NA | NA |
|  |  | Colletidae:lnITD | -0.14 (-0.44 – 0.13) | 1330 | NA | NA |
|  |  | Halictidae:lnITD | 0.02 (-0.15 – 0.18) | 1195 | NA | NA |
|  |  | Megachilidae:lnITD | 0.02 (-0.22 – 0.26) | 1135 | NA | NA |
|  |  | Melittidae:lnITD | -0.01 (-0.21 – 0.09) | 2688 | NA | NA |
| Random | Region |  | 0.29 (0.11 – 0.65) | 1702 | 0.27 (0.1 – 0.64) | 1752 |
|  | Region:Species |  | 0.30 (0.27 – 0.33) | 871 | 0.59 (0.52 – 0.66) | 977 |
|  | Sigma |  | 0.33 (0.32 – 0.34) | 4000 | 0.33 (0.32 – 0.34) | 4000 |

**Table S4B.** Hoverflies: Posterior mean model parameters for best-fitting taxonomic GLMM. Post.: Posterior mean estimate (95% confidence intervals). ESS: Effective sample size. Model formula: ln dry weight ∼ ln ITD + Sex +Subfamily + Sex:lnITD + Subfamily:lnITD + (1|Region/Species).

| *Taxo.* GLMM |  |  |  |  |
| --- | --- | --- | --- | --- |
| Effect type |  |  | Post. | ESS |
| Fixed | Intercept |  | -0.13 (-0.92 – 0.37) | 1287 |
| | $\ln \text{ ITD}$ | | 2.50 (2.23 – 2.76) | 2462 |
|  | Sex | Male | -0.09 (-0.33 – 0.14) | 2811 |
| | | Male:$\ln \text{ ITD}$ | -0.11 (-0.36 – 0.15) | 2885 |
|  | Subfamily | Syrphinae | -0.02 (-0.35 – 0.31) | 2349 |
| | | Syrphinae:$\ln \text{ ITD}$ | -0.16 (-0.48 – 0.18) | 2037 |
| Random | Region |  | 0.31 (0.06 – 0.81) | 1664 |
|  | Region:Species |  | 0.2 (0.12 – 0.28) | 1357 |
|  | Sigma |  | 0.44 (0.41 – 0.48) | 4000 |

**Figure S1.**
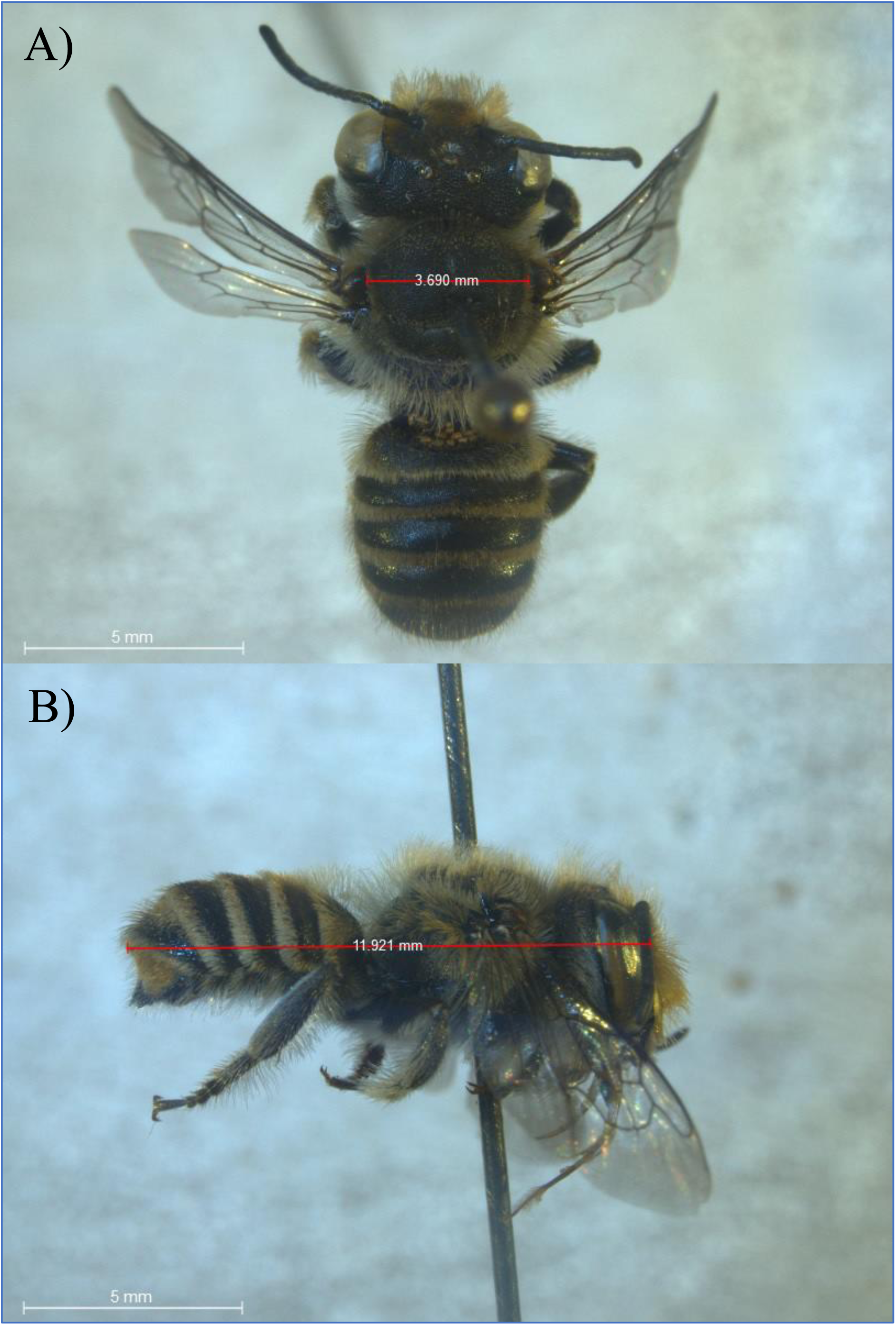
Photographs of A) intertegular distance (ITD) and B) body length (BL) measurements. Specimen is an Australian ♂ *Megachile (Eutricharaea) serricauda*.

**Figure S2.**
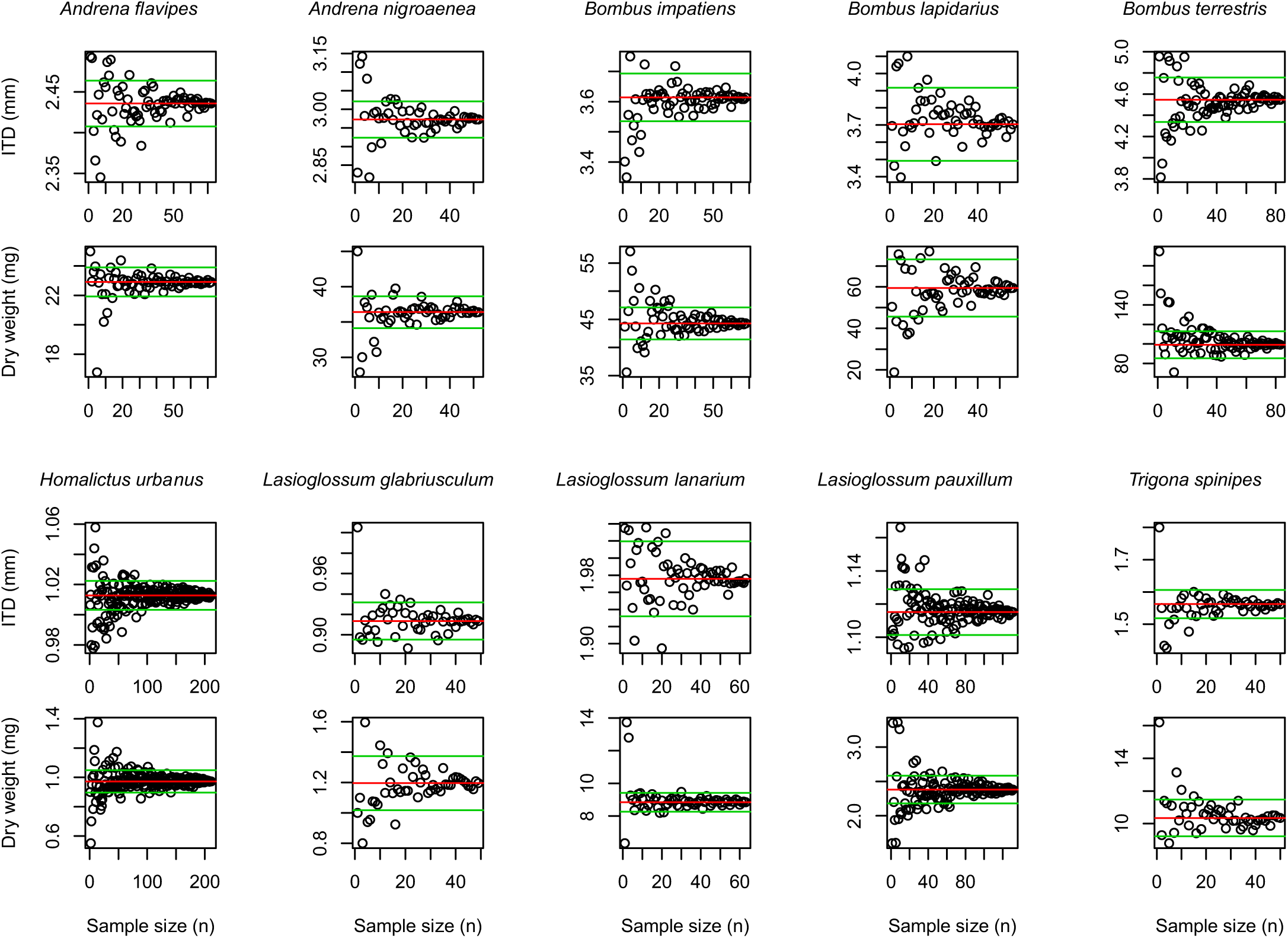
Intraspecific variation in intertegular distance (ITD) and body size (dry weight) in relation to sample size in the 10 most abundant bee species. Red lines denote the total trait mean and green lines represent 95% confidence intervals.

